# AstroLogics: A simulation-based analysis framework for monotonous Boolean model ensemble

**DOI:** 10.1101/2025.11.17.688236

**Authors:** Saran Pankaew, Vincent Noel, Loic Paulevé, Denis Thieffry, Emmanuel Barillot, Laurence Calzone

## Abstract

Boolean networks (BNs) have emerged as versatile tools for modeling cellular regulatory mechanisms due to their ability to capture key biological features despite their simplicity. Multiple BN synthesis methods, which aim to infer BNs with dynamics that corresponding to the experimental data, have emerged in a recent decade. However, these methods often posed a challenge as multiple BN candidates or “model ensemble” can be generated. While these ensembles are valuable for representing cell populations and their heterogeneity, current methods typically treat them as single components without examining their constituent features. We present AstroLogics, a novel framework designed to analyze and identify differences in both dynamical behavior and logical regulation within BN model ensembles. The framework calculates dynamical distances between BNs through exploration of their state transition graphs (STGs), enabling clustering of similarly functioning models that may represent different cellular fates or signaling mechanisms. AstroLogics also identifies key logical properties that govern each cluster, highlighting the core regulatory structures that differentiate model behaviors. Our approach leverages MaBoSS, a stochastic simulation tool that implements the Boolean Kinetic Monte-Carlo algorithm to address time interpretation in BNs. This probabilistic approximation method allows efficient probing of BN dynamics through stochastic simulations, overcoming the computational limitations of exhaustive STG analysis. Our framework also provides powerful visualization and classification capabilities for model ensembles. Through multiple use cases, we demonstrate how AstroLogics facilitates comprehensive analysis of model diversity, and discovery of key regulatory structures within BN ensembles.

## Introduction

Genomic research is increasingly focused on understanding cellular decision-making, the process by which cells integrate diverse signals to control essential functions such as differentiation, proliferation, and response to stress. Gaining insight into these decision-making mechanisms is critical to uncovering the molecular basis of diseases and developing targeted therapeutic strategies. However, because these processes are governed by complex, highly integrated, and parallel networks of interactions, computational models are indispensable tools for their study. At the heart of cellular decision-making lies the gene regulatory network (GRN), a complex system of interactions among genes, transcription factors, and other regulatory molecules that determine when and how genes are expressed.

These networks process both internal and external cues to guide cellular behavior. Due to their complexity, computational modeling is essential to understand their structure and function. Various modeling approaches have been developed for GRNs, including linear models, Bayesian networks, neural networks, differential equations, and stochastic models. Each approach offers different advantages depending on the biological detail required, computational complexity, and available data. Among these, Boolean networks (BNs), first introduced by Kauffman [1] and Thomas [2], have emerged as particularly versatile and intuitive tools for modeling cellular regulatory mechanisms. A BN represents genes and proteins as binary variables (active or inactive) and describes their regulatory relationships using logical functions. Despite their simplicity, BNs can capture key features of biological systems, including feedback loops, attractor states, and robustness to noise. Their discrete nature makes them especially useful in situations where quantitative data is limited and their interpretability facilitates biological insight. As a result, BNs have gained broad adoption in systems biology and have been successfully applied to model processes such as cell fate determination and signal transduction [3, 4, 5].

A recent critical challenge in the field of BNs lies in their inference or synthesis process, which seeks to reconstruct BN models from experimental data and existing biological knowledge. Numerous methods have been developed to infer BNs from time-series data using diverse algorithmic strategies [6, 7, 8, 9]. These approaches typically apply biological constraints to reduce the models’ search space. However, such constraints concern a limited number of conditions or contexts that can lead to multiple candidate models or what is often referred to as “BN model ensemble” or “model ensemble”, in short.

The idea of ensemble modeling has gained momentum with machine learning, notably with gradient-boosted trees and random forests. This approach aims at capturing both model uncertainty and potential model variability across different biological systems (such as different cells, within biological samples). The use of model ensembles spans multiple applications, for instance, ensembles of random BNs have been employed to show emerging properties of families of BNs sharing properties related to their architecture or logic rules [10, 11]. In another instance, BN ensembles were used to assess dynamical properties of qualitative differential equations [12]. Our previous work has demonstrated the use of BN ensembles as single models that represent cell populations and account for their potential heterogeneity [13]. In this work, we constituted ensembles of BNs sampled from the multitude of models compatible with network architecture and predefined dynamical properties. These models were then simulated asynchronously, and the simulations were aggregated through averaging.

However, this method posed a significant limitation, in which the model ensemble is typically treated as single components without thorough examination of the constituent BNs’ features. These features include similarities in BN dynamics and logical functions across the ensemble. Analyzing such characteristics is essential for evaluating the diversity within model ensembles, the different points of dynamical bifurcation in cell fates, and identifying key logical regulations that serve as core regulatory structures.

This challenge revealed a gap in comprehensive tools to analyze and identify differences in both dynamical behavior and logical regulation of BNs in model ensembles. To address this gap, we present AstroLogics, a framework designed for calculating dynamical distances between BNs, clustering similar models, and identifying key logic regulations that govern each cluster. We described the concept of the framework, and illustrated its function through multiple use case examples.

## Design and implementation

### Introduction to Boolean modeling

A Boolean network (BN) consists of a **regulatory graph** (*V, E*) and **dynamical parameters** (*K*). The **regulatory graph** is a signed directed graph where *V* = *{v*_1_, *v*_2_, …*v*_*n*_*}* represents *n* nodes (genes, proteins, complexes, etc.) and *E* represents signed directed edges showing interactions between components.

The **dynamical parameters** are represented by associating each *v*_*i*_ with a discrete variable *D*_*i*_ = 0, 1, determined by logical functions *K* = *{K*_*i*_*}*_*i*=1,…,*n*_. These functions use logical connectors *∨* (AND), *∧* (OR) and *¬* (NOT) to define state transitions based on the regulation of the component states.

Time in Boolean modeling reflects the sequential nature of the state transition. The update mode of these transitions is crucial to capture the dynamics of the systems. The updating methods can be crudely categorized into two main methods: (i) synchronous updating, all components in the network update their states simultaneously, creating a deterministic system where each configuration has exactly one successor. (ii) asynchronous updating allows each individual node to update a each transition, generating all possible elementary transitions and creating a non-deterministic system where configurations can have multiple successors [14].

In this work, we focus on the asynchronous updating method. As the information on the state changes (updating timing) is often unknown, the asynchronous updating scheme offers a more flexible approach to capture overall dynamics.

#### Dynamical properties obtained from Boolean models

Understanding the dynamics of the model requires an understanding of two key concepts: states and transitions. A state of the system is represented by a vector *s* = (*s*_1_, *s*_2_, *s*_3_, …, *s*_*n*_) where *n* denotes the number of components, and *s*_*i*_ the state of component *v*_*i*_. The graph representing the whole solution space, including states and the possible transitions is referred to as the state transition graph (STG), where nodes correspond to states and edges to the possible transitions.

From the STG, multiple features can be extracted including attractors and reachability. The terminal states of the STG are referred to as attractors and defined as state sets with no escape transitions. Attractors come in two types: single attractors (stable states) where component values remain constant, and cyclic attractors which are sets of mutually reachable states with no outward transitions. These attractors have important biological significance. Single-state attractors can represent cell differentiation states [5] or final cellular fates (proliferation, death, senescent states, etc.) [15], while cyclic attractors typically represent periodic behaviors like cell cycles or circadian rhythms [16].

When analyzing a model, we characterize its main dynamic properties, including attractors and their accessibility. While small models allow straightforward attractor identification through STG exploration, larger models with complex interactions face combinatorial explosion. This necessitates alternative methods to capture the crucial information from the BN’s dynamics, such as attractors, reachability, basin of attraction or trap spaces.

### Stochastic simulations of Boolean models via MaBoSS

Computing STG transitions requires considerable computational power, and is often hard to visualized in large model. To address this, a simulation based approach allows an approximation of the STG through stochastic simulation. In this work, we utilized MaBoSS, a stochastic simulation tool for continuous-time Markov processes in C++ [17, 18]. It implements a Boolean Kinetic Monte-Carlo (BKMC) algorithm, assigning transition rates to node activation and deactivation to model different timescales and introduce continuous time in a Boolean framework. The tool performs multiple random walks on the probabilistic STG, calculating the activation likelihood for each node and state. Users can modify parameters, such as initial conditions and node activity status for thorough analysis. The simulation produces a vector of activation probability for states in the STG, which can be converted into activation probabilities for each node in the model.

### AstroLogics framework

The aim of AstroLogics framework is to provide a tool for analyzing and identifying differences between Boolean models within the model ensemble. We focus on two major properties of each BN: dynamical and logic properties, where differences between BNs are determined by their dynamical properties which is governed by logical properties (logical function *K*, as mentioned previously). Thus, we developed a framework which seeks to determine the distance between BNs through the exploration of BN’s STG and determine distances between each model. The distances between BNs in the cohorts enable us to perform clustering to identify groups of similarly functioning models, which could be associated with different cellular fate, or signaling mechanism. Finally, we can identify key logical properties, which consist in highlighting differences in the logical equations that separate the clusters. We illustrate the pipeline of this framework in Figure 1.

**Fig. 1.**
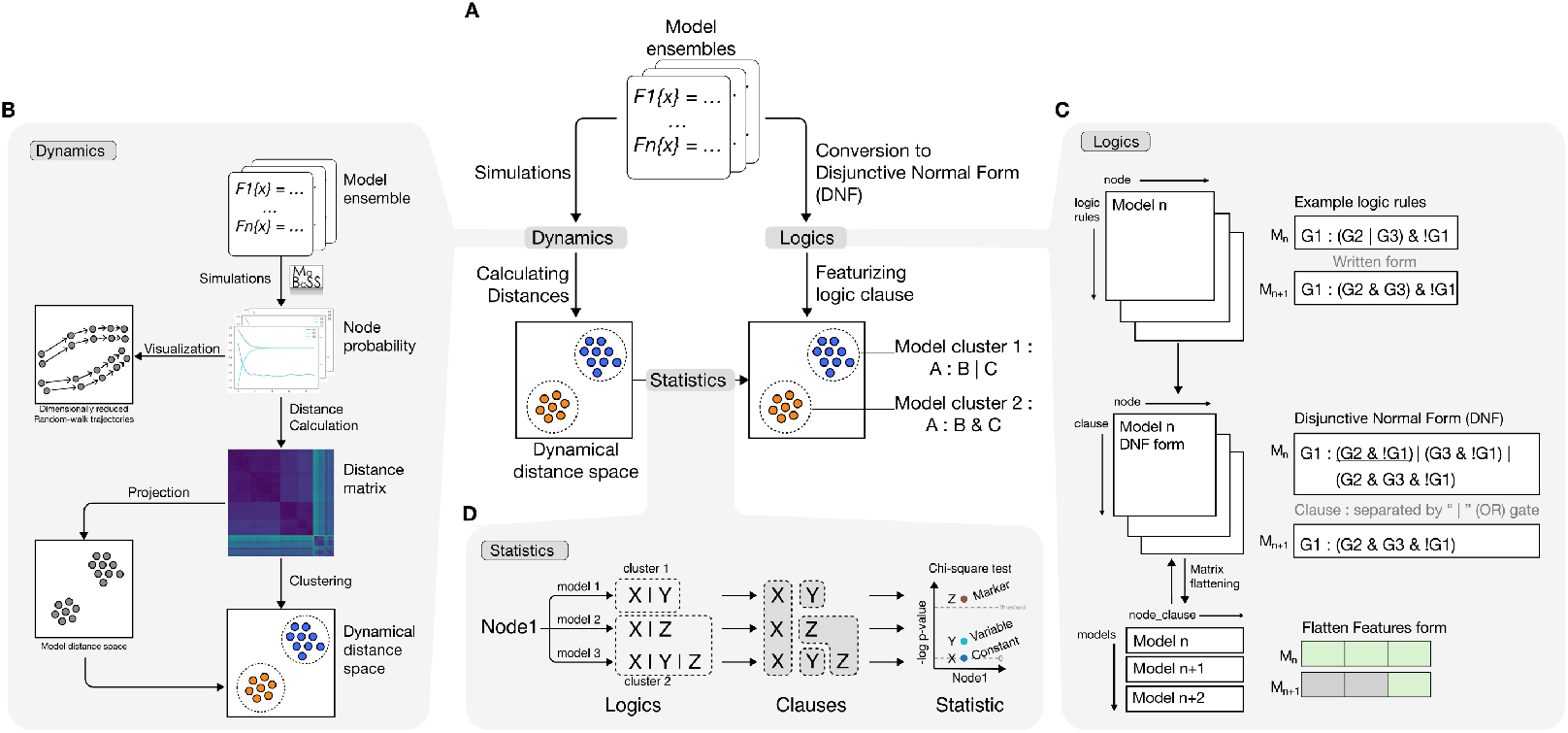
Overview of AstroLogics framework. (A) Overview of the pipeline which is separated into two major parts according to BN properties. (B) The pipeline for comparing BN dynamical properties via two different approaches (Succession Diagram and MaBoSS simulation). (C) The processing pipeline of each BNs’ logic property. (D) The pipeline for performing statistical test of logic clauses between model clusters.

#### Dynamical features

As previously mentioned, STG calculation requires intensive computational power. Therefore, different methods have been developed to comprehend key STG features. While attractors indicate model dynamics, networks with identical attractors may exhibit different dynamical properties based on their STG structure. With MaBoSS, we specify initial parameters for each model ensemble: initial states, simulation time steps, and random-walk sample numbers. By default, our pipeline uses random initial states for full state space exploration, though specific initial states can be specified for detailed subsection analysis. The simulation produces activation probabilities for each state, which can be converted into node activation probabilities and stored as a time series matrix. From this matrix, we calculate model differences using either terminal-timepoint data (Euclidean distance) or full trajectory data via Dynamic Time Warping (DTW) [19]. We employ Non-metric Multidimensional Scaling (NMDS) [20, 21] to visualize the distance matrix in 2-dimensional space, then perform k-mean clustering to identify BN clusters with similar functions and project them onto the NMDS space (Figure 1B).

#### Logic features

Boolean functions describe how each node gets activated using logical operators like *∧, ∨*, and *¬*. Since comparing these strings systematically is challenging, we convert them into features matrices. We developed a pipeline to transform Boolean equations into comparable features matrices (Figure 1C). First, we convert each model’s logical equations into Disjunctive Normal Form (DNF), which represents Boolean expressions using OR-connected “clauses” where each clause combines variables using AND. DNF’s canonical nature allows any Boolean expression to be converted, and simplifies Boolean equation complexity [22]. For each Boolean network, we treat individual component clauses as features, creating a matrix where columns represent nodes, rows represent possible clauses, and values indicate presence of each node-clause logic. We flatten this matrix into a vector of ‘*node*_*clause*’ for each model, then combine these vectors to create a final matrix where columns represent *node*_*clause* and rows represent individual models.

#### Linking Dynamical features with Logical features

After identifying clusters of Boolean Networks (BNs) based on their dynamics property, we sought to understand the underlying logic structure differentiating these clusters by linking dynamical and logic features (Figure 1D). To do so, we utilized a statistical method, similar to marker finding of single-cell analysis, to identify key logic features for each model clusters. We employed Chi-square statistical tests on each logic feature to determine its association with specific model clusters. Based on p-values and a defined threshold, we categorized logic features into three groups: constant features (present in all models), variable features (high p-value), and marker features (low p-value, statistically significant between clusters). Results can be visualized through Manhattan plots, summary bar plots, or heatmaps illustrating logic clause distribution across models. This approach identified key logic properties distinguishing different BN clusters, providing insights into which regulatory mechanisms are responsible for specific dynamical behaviors.

## Results

### Demonstration of AstroLogics functions

To demonstrate the function of AstroLogics, we first utilized a network of Central Nervous System (CNS) differentiation model [23] (Figure 2A), which has been used as a tutorial network for BoNesis [24]. Following the tutorial, we generated an ensemble of 88 models https://bnediction.github.io/bonesis/tutorials/tour.html. Applying our approach, we first looked at diverse groups of attractors that each BN could reach. We found that within the model ensembles, BNs can be separated into 2 major groups according to the set of reachable attractors (Figure 2B). We then asked what are the key logic properties that distinguished these two groups of BNs. Following the logical comparison pipeline of AstroLogics (subsection AstroLogics framework in **Design and Implementation**), we were able to identify 3 major groups of logic features: I). constant features, which are the key logic regulations that are included in all BNs and can be seen as the backbone of the model, II). variable features, which are the logic regulations that may vary between BNs but are not statistically significant between clusters of BNs, and III). marker features, which are the logic regulations that are statistically significant between two clusters of model. To identify the nodes of interest, we created a bar plot showing the number of possible logic variations for each node, and the percentage of logic clause categories for each node (Figure 2C).

**Fig. 2.**
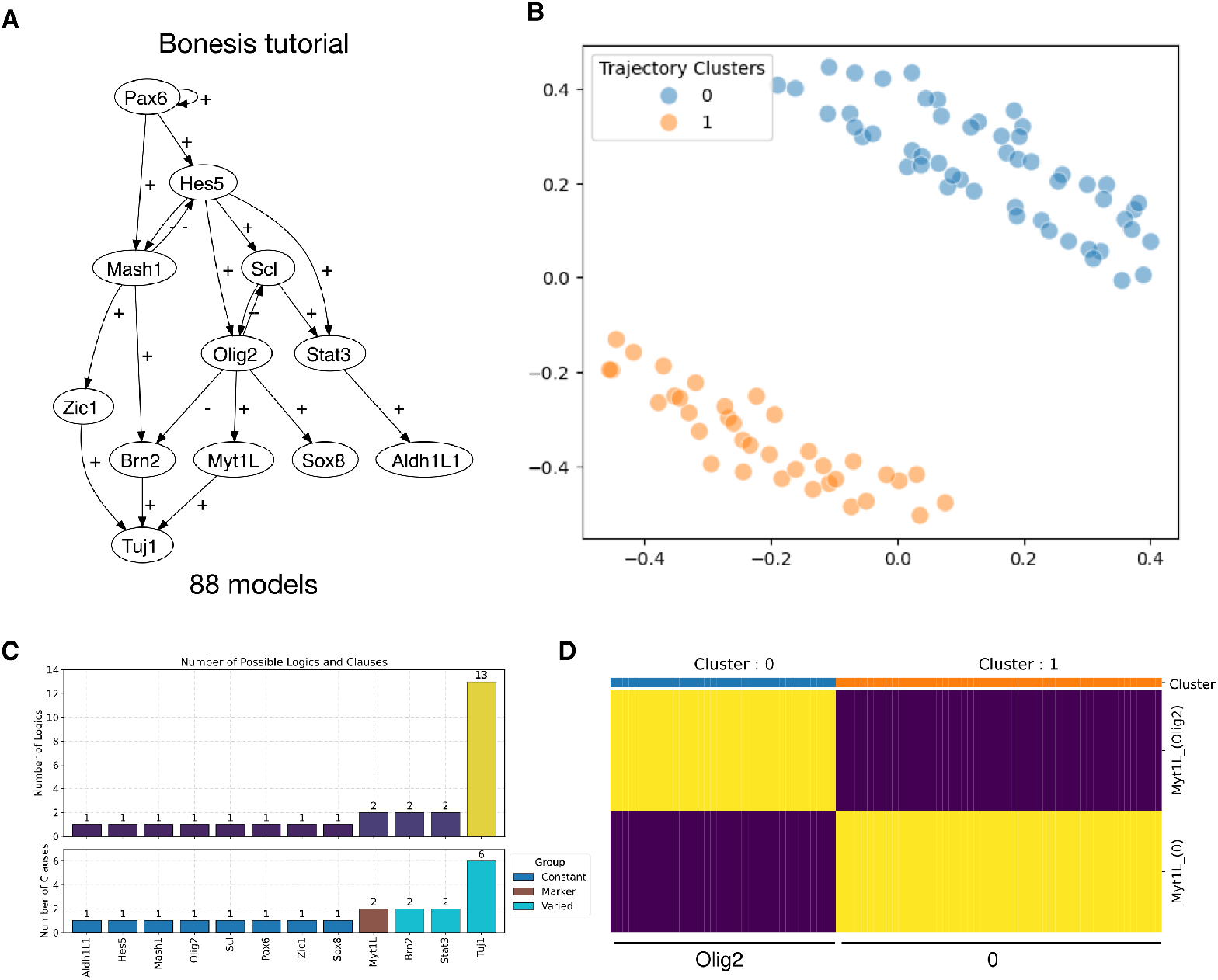
Example of AstroLogics framework utility. (A) An example network of central nervous system (CNS) differentiation (B) Distance of models calculated by differences in the MaBoSS trajectories, visualized using MDS. (C) Barplot of number of clauses for each node and percentage barplot represents proportion of logic features in each category (Dark blue = Marker feature, Brown = Constant feature, and light blue = Variable feature). (D) Heatmap showing the different logic features of node Myt1L. Each row represents the Myt1L logic feature, while each column represents a model in the model ensemble, with column color indicated by the cluster of the model.

From this visualization, we found that 8 nodes contain only one clause (Adh1L1, Hes5, Mash1, Olig2, Scl, Pax6, Zic1 and Sox8). We found that Myt1L, Brn2, Stat3, and Tuj1 nodes contain multiple logic equation forms, resulting in differences between models. However, of all these nodes, we found that only Myt1L contains logic features that are categorized as marker features. From this visualization, we concluded that Myt1L was a key node, in which the logic features are the key identifier between two groups of models. We then displayed the different Myt1L logic features in the form of heatmap (Figure 2D), with color label indicating which cluster each model belongs to. From this visualization, we found that in cluster 0, Myt1L is regulated by Olig2 while in cluster 1, Myt1L is deactivated (0). More analysis can be done on the possible interpretations of these two clusters of models led by Myt1L regulation, and if they can be associated with two differentiated states or cell fates.

This analysis could be achieved by few lines of code. We included the example script to replicates all plots shown in this figure in Supplementary Figure S1, as well as detailed tutorials on our github page https://github.com/sysbio-curie/AstroLogics.

### Comparison between trajectory clustering and attractor analysis approach

Inspired by the AEON approach [25], which utilized long-term behavior of BN to characterize BNs into groups, we implemented an attractor calculation function via `boolsim` [26] to characterize models based on reachable attractors, which were then used to group models into clusters. We compared this attractor-based clustering with our simulation-based approach using 19 selected networks from Kadelka et al., 2024 [27]. For each network, we generated 1000 random BN model ensembles using Bonesis. We calculated attractor states for each model and classified them into clusters. In parallel, we performed MaBoSS simulations on all models, obtained node activation probabilities at the endpoint, and calculated Euclidean distances between models. Since the optimal number of clusters for simulation-based clustering was unknown, we performed k-means clustering with varying cluster numbers from 1 to n (where n is the number of attractor-based clusters) and calculated the Adjusted Rand Index (ARI) for each to compare clustering similarity. We selected the maximum ARI for each network, as using n clusters in simulation-based clustering does not guarantee maximum agreement with attractor-based results.

We showed the results in the Table 1, which shows the list of 19 selected models, along with the characteristic of the network, including number of nodes and interactions, along with the number of model generated from Bonesis. We listed the number of clusters based on the attractor identification approach, along with the ARI-score from the terminal-timepoint simulation. We found that several model shows almost complete agreement with the attractor identification approach, with 16 out of 19 models shows ARI score >0.85, while the remaining 3 models shows low ARI score (<0.45). We further investigated this issue in supplementary analysis 1. From the results, we found that by using the states activation probability, we can define reachable attractors for each BN similar to the attractor calculation method. However, due to the nature of probabilistic approximation, different probability are assigned to each attractor, depending on the level of states reachability. We found that in some attractors, the probability of reaching these states are minimal, resulting in minimal distances between model with and without these attractors. Moreover, we showed that the conversion between states activation probability to node activation probability resulted in loss of information, which further lower agreement between attractor-based and simulation-based clustering (Supplementary Analysis 1).

**Table 1.** Correspondent between attractors and simulation based clustering.

| project name | Nodes | Interactions | num model | num attractors groups | Max ARI (endpoint) |
| --- | --- | --- | --- | --- | --- |
| Body Segmentation in Drosophila 2013_23520449 | 14 | 12 | 1000 | 86 | 1 |
| Death Receptor Signaling_20221256 | 25 | 45 | 1000 | 16 | 1 |
| FGF pathway of Drosophila Signaling Pathways_23868318 | 14 | 24 | 1000 | 570 | 1 |
| HH Pathway of Drosophila Signaling Pathways_23868318 | 11 | 30 | 1000 | 1000 | 1 |
| Mammalian Cell Cycle_19118495 | 19 | 44 | 1000 | 7 | 1 |
| Processing of Spz Network from the Drosophila Signaling Pathway_23868318 | 18 | 20 | 1000 | 152 | 1 |
| Toll Pathway of Drosophila Signaling Pathway_23868318 | 9 | 11 | 16 | 6 | 1 |
| VEGF Pathway of Drosophila Signaling Pathway_23868318 | 10 | 18 | 1000 | 390 | 1 |
| Regulation of the L-arabinose operon of Escherichia coli_28639170 | 9 | 16 | 1000 | 130 | 0.990976 |
| SKBR3 Breast Cell Line Long-term ErbB Network_24970389 | 21 | 52 | 1000 | 41 | 0.977437 |
| Cardiac development_23056457 | 14 | 24 | 1000 | 6 | 0.977327 |
| HCC1954 Breast Cell Line Short-term ErbB Network_24970389 | 11 | 33 | 1000 | 304 | 0.962528 |
| Lac Operon_21563979 | 10 | 22 | 1000 | 61 | 0.961812 |
| SKBR3 Breast Cell Line Short-term ErbB Network_24970389 | 11 | 29 | 1000 | 268 | 0.931461 |
| B cell differentiation_26751566 | 17 | 38 | 1000 | 287 | 0.907193 |
| BT474 Breast Cell Line Long-term ErbB Network_24970389 | 21 | 41 | 1000 | 249 | 0.887925 |
| Human Gonadal Sex Determination_26573569 | 19 | 76 | 1000 | 6 | 0.442307 |
| HCC1954 Breast Cell Line Long-term ErbB Network_24970389 | 21 | 42 | 1000 | 542 | 0.362385 |
| Cortical Area Development_20862356 | 5 | 11 | 1000 | 5 | 0.138381 |

### Using transient dynamics of BNs to cluster models

In addition to terminal-timepoint simulation, MaBoSS simulation provides node activation probabilities throughout simulation time-points, enabling analysis of transient dynamics. While attractor states are valuable for identifying long-term system behaviors corresponding to phenotypes or cell fates, they offer only a limited view of model dynamics. The path a system takes before reaching an attractor often contains crucial biological information, such as transient gene activation and signaling cascades. This is particularly important in developmental and immune systems, where the timing and order of activation play critical roles beyond the final state. Furthermore, two models with identical attractors may follow significantly different paths to reach them, which impacts both interpretability and potential intervention strategies.

To utilize this information, we keep the whole simulation trajectory from MaBoSS (all nodes activation probability at all simulation time-point). We apply the Dynamic Time Warping (DTW) algorithm, implemented in the tslearn package to calculate the distance between the simulation trajectories of each models. This resulting in the distance matrix similar to the Euclidean distance matrix, that could be used to visualize in the MDS latent space, and/or perform clustering to identify model clusters.

After implementing this method, we first verify whether this approach changes the performance of model classification according to different attractor groups. To answer this question, we undertook the same comparative evaluation as the previous section. We re-calculate models distances using the dtw method and perform k-mean cluster, varying the number of clusters from 1 to n, where n is the number of clusters obtained from the attractor states classification. We then calculated the similarity of clusters with the attractor-based clusters, using ARI and selected the maximum value for each network. We found that there is no observable differences in the ARI score between the two approach (terminal-timepoint or whole trajectory), suggesting that despite the including transient dynamics our approach still able to classify models according to their reachable attractors without compromising the efficacy (Supplementary Figure S2).

We then seek to demonstrate when the whole trajectory dynamics may add some additional insight. To do so, we examined a model ensemble from the Herault hematopoiesis study (n = 616) [28] where all models contains only one attractor group (Figure 3A). We performed MaBoSS simulation in all 616 models, and utilized terminal-timepoint simulation data to calculate model distance and project onto NMDS space. We found that the distance between BNs show no clear clustering within the model ensemble, reflecting the single attractor groups of the model. However, upon utilizing the data of the whole simulation trajectory we found a clear separation of BNs into two groups (Figure 3B). Following the AstroLogics pipeline we performed statistical analysis to identify nodes which are statistically different between two model clusters. We identified CDK46CycD as a marker node (Figure 3C). To look into the differences in this node’s dynamics between two BNs group we plotted the nodes activation probably of all BNs separated by their clusters. The result showed subtle early-stage differences that eventually converged to zero at steady state (Figure 3D). Finally, to identify the key differences in the logic clause, we visualize the differences in the logic clause of CDK46CycD between two BNs clusters using heatmap (Figure 3E). The results revealed that cluster 0 models used the logic (Bclaf1 *∧* Myc), while cluster 1 models employed Bclaf1 *∨* Myc.

**Fig. 3.**
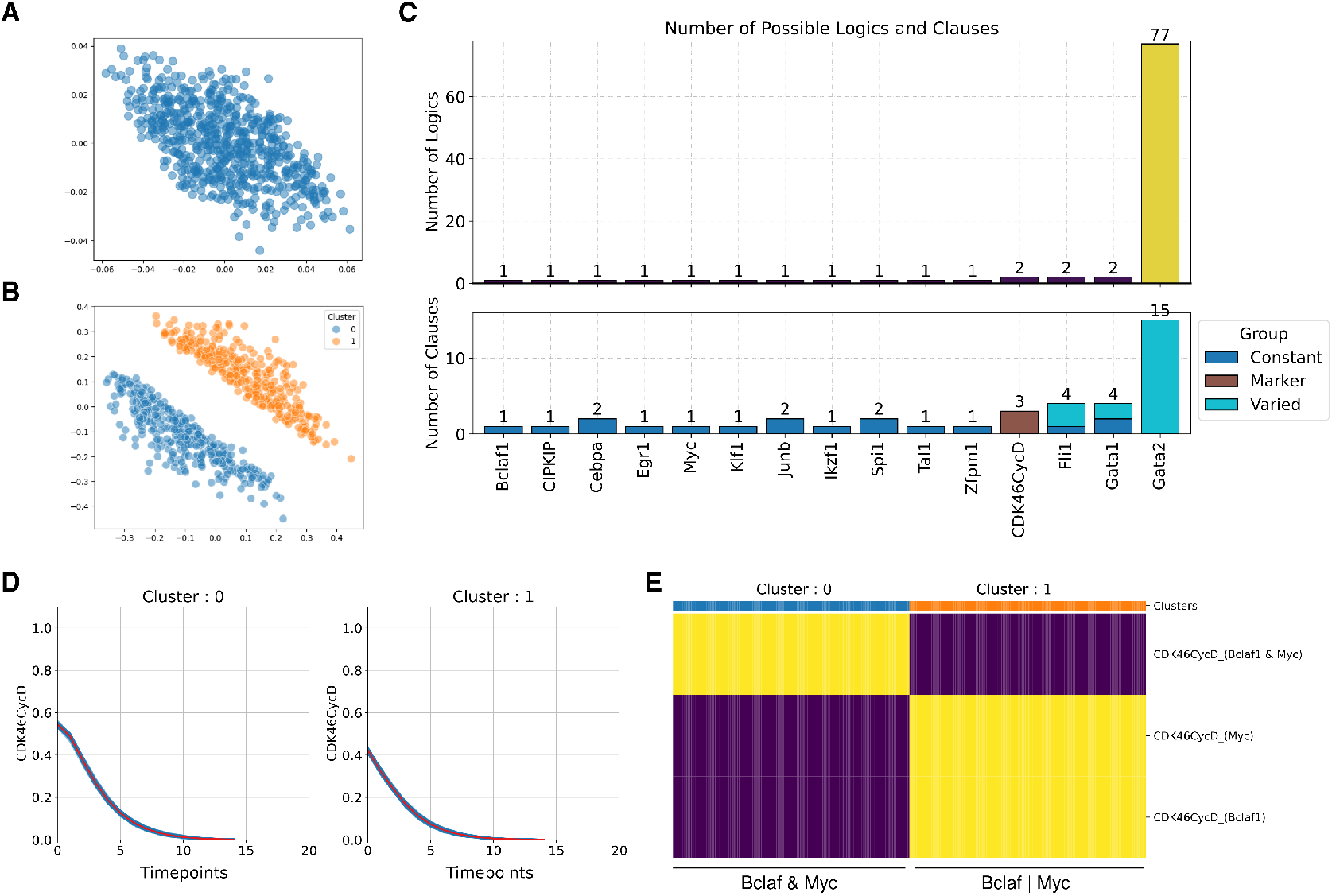
Analysis of transient dynamics in Boolean network models. MDS projection of BNs distances from Herault hematopoiesis model ensemble (n=616) where (A) utilized terminal-timepoint of the simulation data and (B) utilized the whole trajectory from the simulation data. (C) Barplot of number of logical variation for each node (upper), and a barplot represents proportion of logic features in each category (Dark blue = Marker feature, Brown = Constant feature, and light blue = Variable feature) (lower). (D) Visualization of CDK46CycD node activation probability at all simulation simulation timesteps (E) Heatmap shows the different logic features of node CDK6CycD. Each row represents the CDK6CycD logic feature, while each column represents a model in the model ensemble, with column color indicated by the cluster of the model.

### Case study: cancer invasion model

Another major advantage of our simulation framework lies in its ability to approximate model dynamics. This approximation significantly reduces computational time, particularly when dealing with complex models that contain numerous attractor states and would otherwise require extensive computational resources. To demonstrate these capabilities, we utilized a cancer invasion model described in Chevalier et al., 2020 [13] (Figure 4A). We compared the computational time required for attractor calculation versus simulation time at varying sample counts (1000, 2500, 5000, 100000). Our findings revealed that regardless of sample count, simulation via MaBoSS demonstrated significant time advantages compared to traditional attractor identification methods (via boolsim implementation)(Figure 4B). To further illustrate our framework’s utility, we followed the methodology and constraints outlined in [13] to generate a model ensemble of 1024 models. We performed MaBoSS simulations on these models and calculated distances between models. By projecting these distances onto the MDS latent space and applying k-means clustering, we identified five distinct model clusters, as shown in Figure 4C. Analysis of these clusters revealed two major groups (clusters 0 and 2) within the model ensemble (shown in dotted line). Examination of logic features using heatmap, we found similar profiles between these two clusters compared to others (Supplementary figure S3). This similarity suggests that these clusters maintain stable sets of logic features, reflected in reduced logic variation across the model and fewer nodes characterized as varied or marker nodes (Figure 4D). To identify key dynamical differences between these two model clusters, we first determined the nodes showing the most dynamical variation during simulation across all models. We selected nodes which contains marker clauses which includes CDH1, EMT, Invasion, Apoptosis, Migration, SNAI2, GF, AKT1, AKT2, ERK and ZEB1. We visualized node activation trajectories of these selected nodes. We found a major differences in GF activity between clusters 0 and 2 (Figure 4E) while other node exhibits minor difference in the trajectory (Supplementary figure S4). Further investigation of logic features between models in these clusters revealed distinct sets of GF logic equations as shown in Figure 4F. These two different dynamics suggested the two groups of logical function where GF is or is not subjected to a negative feedback, allowing the users to compare with the actual biological knowledge/data and subset groups of models.

**Fig. 4.**
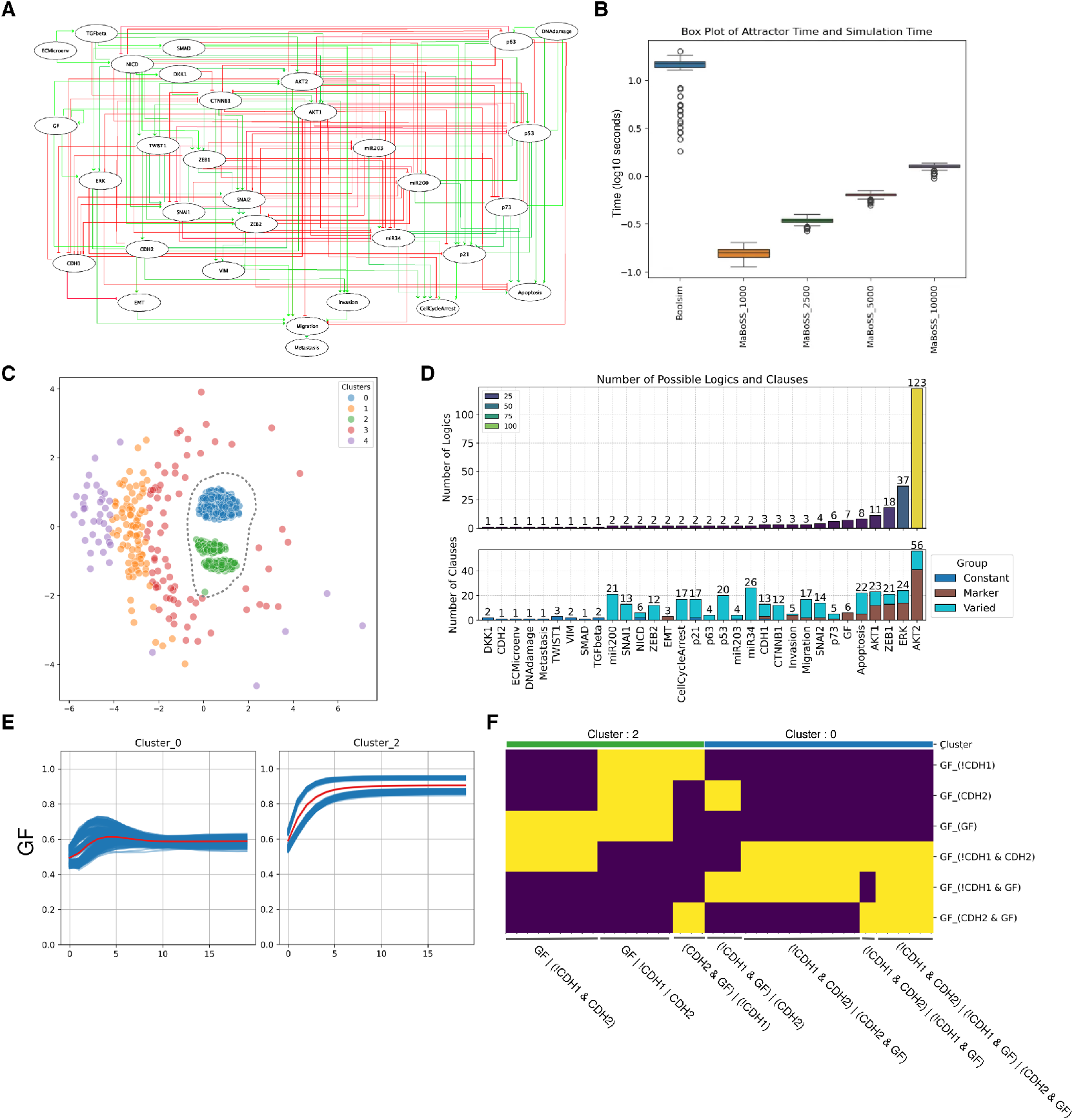
Applying AstroLogics to analyze a complex cancer invasion model. (A) Network diagram of the cancer invasion model [13] showing key nodes and interactions involved in the EMT process of breast cancer. (B) Computational time comparison between traditional attractor calculation methods and MaBoSS simulation at varying sample counts (1000, 2500, 5000, 100000). The y-axis represents log10 second. (C) MDS projection of model distances revealing six distinct clusters from 1024 models generated following Chevalier et al.’s methodology and constraints. The selected cluster 0 and cluster 2 are indicated by dashed line. (D) Comparison of logic variation between the two predominant model clusters (0 and 2), showing number of logical equation variance (upper) and number of possible clauses (lower) for each node, while the color of the stacked bar plot represents the proportion of clause that are categorized as constant (dark blue), varied (light blue) and marker (brown) clauses. (E) Visualization of node activation probabilities for GF node in clusters 0 and 2, highlighting substantial differences in GF activity patterns. (F) Heatmap shows the different logic features of node GF. Each row represents the GF logic feature, while each column represents a model in the model ensemble, with column color indicated by the cluster of the model.

## Discussion

The field of Gene Regulatory Network (GRN) inference has advanced significantly over the past five years, with many new methodologies emerging [6, 7, 8, 9]. This progress has generated BNs model ensembles that can be refined by applying additional criteria to narrow the solution space. Beyond simply restricting options, analyzing these model ensembles provides valuable insights into key inferred information and highlights similarities across models within the ensemble. However, there is a gap in the development of the tools to analyze models within the solution space, therefore we aim to address this gap by presenting AstroLogics framework. AstroLogics is a comprehensive framework designed for analyzing BNs within the model ensemble that arise from gene regulatory network inference methods. It evaluates BN models through two complementary approaches; dynamical properties analysis and logical function evaluation.

In our dynamical properties analysis, we focus on both the long-term behavior of the Boolean Network (BN) and its transient dynamics. Other model classification methods such as AEON and BNclassifier [25, 29] primarily concentrate on classifying BN’s attractors to identify their groups. These methods are adapted to analyze Parameterized Boolean network (PBN), a Boolean network where some of the logical update functions contain parameters (uninterpreted Boolean functions) instead of being fully specified. These methods utilize long-term behavior of the model, or ‘attractors landscape’ to analyze how changes in logical parameters affect the attractors landscape. These tools rely on symbolic algorithms to compute and categorize attractors into three types: stability (single state), oscillation (periodic cycle), and disorder (chaotic behavior), enabling classification of Boolean network models based on their long-term dynamical properties. However, this tool relies on coarse 3 attractor categories to characterize BNs, and the utilization also requires steep learning curve.

While long-term behavior of the BN can be used as their classifier, its transient dynamic also plays important role in determining the timing and order of activation of signaling cascade of the BNs. These information plays a major part in the interpretation in the real biological behavior and the potential intervention strategy. We compared our simulation approach with the attractor identification approach to evaluate different methods of analyzing dynamics. Although our framework shows high correspondence with the attractor identification method in most selected models, we observed discrepancies in some networks. These discrepancies result from probabilistic approximation by MaBoSS, which convert the asynchronous BN dynamics in to continuous-time stochastic dynamics. From the Supplementary Analysis 1, we found that in the probability of reaching certain attractor states are minimal, resulting in minimal distances between models with and without these attractors. Despite this discrepancy, we believe that taking into account this probability of reaching the attractor states, will better reflect the dynamics of the BN.

Despite this discrepancy, our simulation approach significantly reduces calculation time when analyzing models with multiple and complex attractor states [Figure 4B]. We therefore present this method as an efficient approximation for exploring and comparing model dynamics. Furthermore, a recent implementation of MaBoSS GPU [30] could potentially scale simulations up to >100 times faster (according to the real-world example models), enabling the simulation of much larger and more complex models [31].

In this work, one major process is the featurization of the logical functions. Methods such as truth tables, K-maps, Logic Circuit Diagrams, and Binary Decision Diagrams (BDDs) provide intuitive representations of logical equations, however, these methods suffer from exponential size explosion as the number of components in the BN increases.The Disjunctive Normal Form (DNF) is a widely used standardized format [32] that simplifies logical equations without complex nesting and facilitates query processing, enabling Boolean function simplification [22]. Leveraging the DNF format, we converted logic clauses into binary features, which allows us to get the overview visualization logical functions via heatmap (Supplementary Figure S3).

Despite its advantages, the DNF format is non-unique, allowing the same logical equation to take multiple representations. Alternatively, a normal format called completed DNF (also known as Blake’s Canonical Form)[33] is the disjunction of all prime implicants, minimal partial assignments that make the function true. The use of this normal form could potentially help reduce the size of the logical equation matrix used for comparison and, more importantly, help identify redundant BNs within the model ensemble. This could serve as a pre-processing step before analyzing BN dynamics.

In our AstroLogics framework, we have integrated method to compare long-term behaviour of BNs using attractors grouping, through boolsim implementation. In recent years, there has been a great interesting in developing a method to identify attractors (both at single-state or cyclical), using different methods such as, symbolic-base, structure-based or decomposition-based (reviewed in [34]), where they expand the computational capacity to larger-scale model, with lower computational time. As the field continues to evolve, it would be interesting to explore and implement new approaches in future versions of our package. These enhancements would further strengthen the capability of AstroLogics to explore complex biological networks while maintaining its user-friendly interface.

Recent advancements in Boolean network analysis have expanded our understanding of network dynamics beyond traditional attractor identification. The concept of trap spaces, regions of state space from which trajectories cannot escape, has emerged as a powerful analytical tool [35]. Within this framework, stable motifs represent self-sustaining components of the network that, once activated, remain fixed regardless of other node states [36]. These stable motifs effectively constrain the system’s behavior by reducing the accessible state space, channeling dynamics toward specific attractors.

These two concept has paved way to novel approach of capture the key structure of the STG (key transition states + long-term behavior) that is not computational intensive allows for comparison across model cohorts. A recent framework developed by Rozum et al., 2021 [37] utilized parity and time reversal transformation to identify stable motifs and their subsequent long-term behavior (attractors). The results of these calculations can be visualized through a Succession Diagram (SD), a directed acyclic graph that represent possible sequences of stable motifs and activation.

With this information in mind, we therefore, have designed our framework with the emphasis of modularity, supporting different model dynamics calculation strategies, including attractor analysis and succession diagrams as implemented by `pystablemotifs` [38]. Our methodology prioritizes interoperability between computational platforms by adhering to the standardized Colomoto tool-suite format. By utilizing established tools from the Colomoto consortium, including `minibn` for model import and boolsim for attractor calculation, AstroLogics ensures seamless integration with existing Boolean modeling workflows. This compliance with Colomoto standards enables researchers to incorporate our framework into their existing analytical pipelines without format conversion requirements. The framework’s modular design supports multiple Colomoto-compatible tools, facilitating cross-platform compatibility and reproducible analyses across different computational environments [39].

## Supporting information

Supplementary Figure, Analysis

## Code availability

The developed AstroLogics package is available on the GitHub https://github.com/sysbio-curie/AstroLogics, along with the code to reproduce figures and table described in the article.

## Author contributions statement

S.P. and L.C. conceived the project. S.P. designed and conducted experiments. S.P. and V.N. worked on the script, packaging, and documentations. L.C. and D.T. consulted the theory of the project. S.P. wrote the manuscript. All authors reviewed the manuscript.

## Acknowledgment

The authors thank the anonymous reviewers for their valuable suggestions. This work was supported by a government grant managed by the Agence Nationale de la Recherche under the France 2030 program, with the reference numbers ANR-24-EXCI-0001, ANR-24-EXCI-0002, ANR-24-EXCI-0003, ANR-24-EXCI-0004, ANR-24-EXCI-0005. L.C. and S.P. were partially supported by Certainty project which is part of the European Union’s Horizon Europe research and innovation program under grant agreement n101136379. L.P. was partly funded by France 2030 project “AI4scMED” operated by ANR (grant number ANR-22-PESN-0002). The authors would like to thank Claudine Chaouiya and Élisabeth Remy from Institut de Mathématiques de Marseille (I2M) for a fruitful discussion.

