## Supplementary Figure, Analysis for "AstroLogics: A simulation-based analysis framework for monotonous Boolean model ensemble"

### Supplementary Information - LogicsEnsemble: An approximate analysis framework for monotonous Boolean model ensemble

Pankaew et al.

November 13, 2025

### 1 Supplementary Data

#### 1.1 Supplementary Figure S1

```python
import astrologics as ast

# Load the model path and create LogicEnsemble object
model_path = '../models/test_bonesis/'
model = ast.ensemble(model_path, project_name = 'test_bonesis')
model.create_simulation()

# Configure simulation parameters
model.simulation.update_parameters(max_time = 15, sample_count = 1000, thread_count = 15)
model.simulation.run_simulation()

# Create trajectory submodule and calculate models distances
model.create_trajectory()
model.trajectory.calculate_distancematrix(mode = 'endpoint')

# Perform MDS (Multidimensional Scaling) for visualization
model.trajectory.calculate_MDS()
model.trajectory.plot_MDS(s = 100, fig_size = (8,8))

# Calculate mode clusters
model.trajectory.calculate_kmean_cluster(n_cluster = 2)
model.trajectory.plot_MDS(s = 100, fig_size = (8,8),plot_cluster = True)

# Create logic submodule and analyze logic rules
model.create_logic()
model.logic.create_flattend_logic_clause()

# Calculate statistics between 2 model cluters and plot summary
model.logic.map_model_clusters(model.trajectory.cluster_dict)
model.logic.calculate_logic_statistic(pval_threshold = 0.0001)
model.logic.plot_logicstat_summary()

# Plot the heatmap of 'MytL1' logic rule
model.logic.plot_logic_heatmap(node = ['MytL1'], figsize = (10,8))
```

**Supplementary Figure S1.** An example script of AstroLogics pipeline to replicate all plots from Figure 2

#### 1.2 Supplementary Figure S2

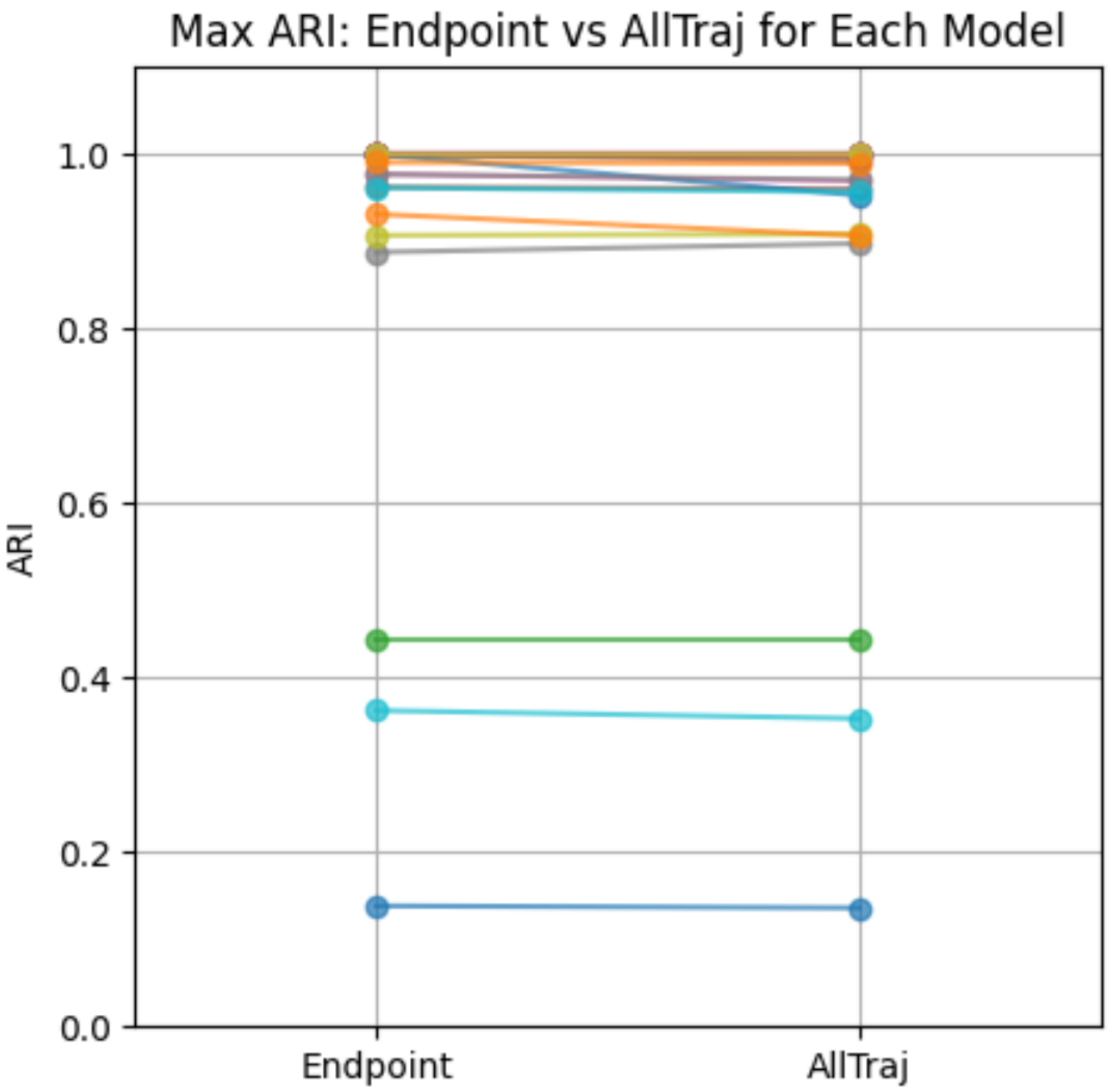


**Supplementary Figure S2.** A comparison of two simulation-based clustering method, in comparison with attractor group clustering. The y-axis shows the correspondant between the attractor group clustering and the simulation based clustering, measured by Adjusted Rand-Index (ARI) with 1 indicates perfect correspondant, and 0 indicates the random agreement. The x-axis shows two simulation-based clustering method: endpoint, which uses the simulation data at the endpoint of the simulation and AllTraj, which uses the whole trajectory of simulation to calculate the distance between models.

#### 1.3 Supplementary Figure S3

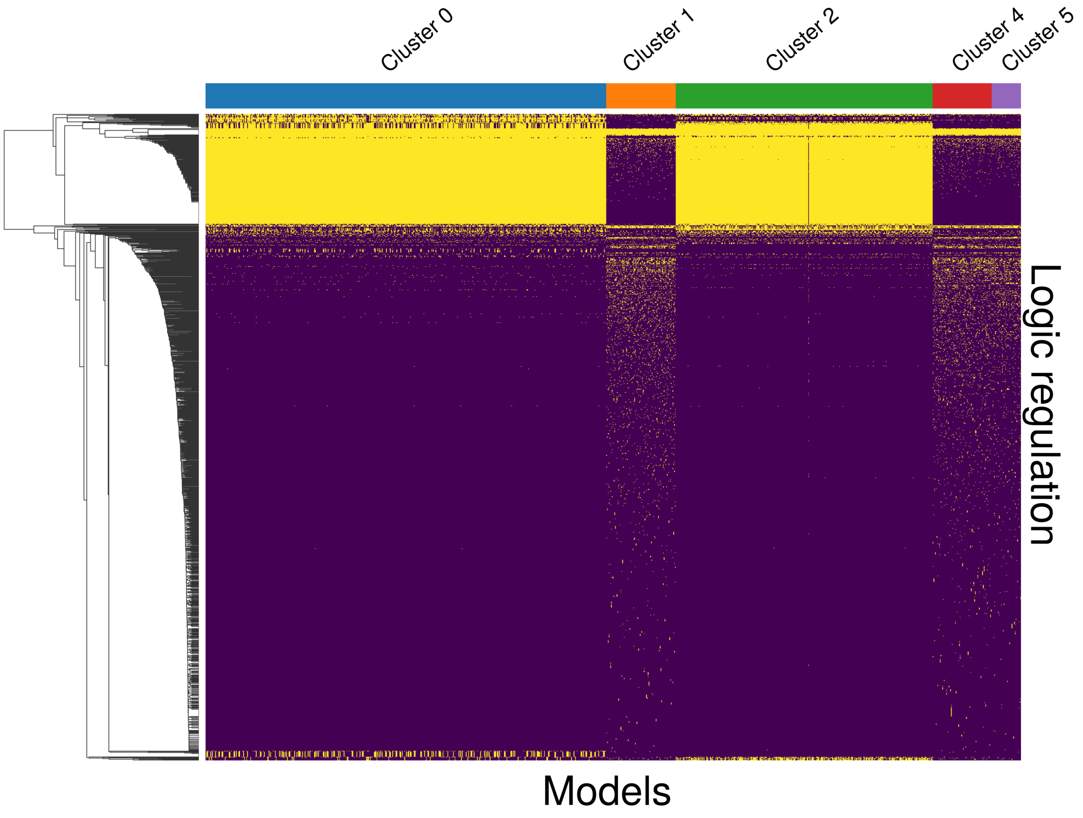


**Supplementary Figure S3.** Heatmap shows different logic features (clauses) (row) in each BN (columns) where yellow color indicates that the BN contains certain logic clause. The color bar on the top of the plot represents clusters where each BN belongs.

#### 1.4 Supplementary Figure S4

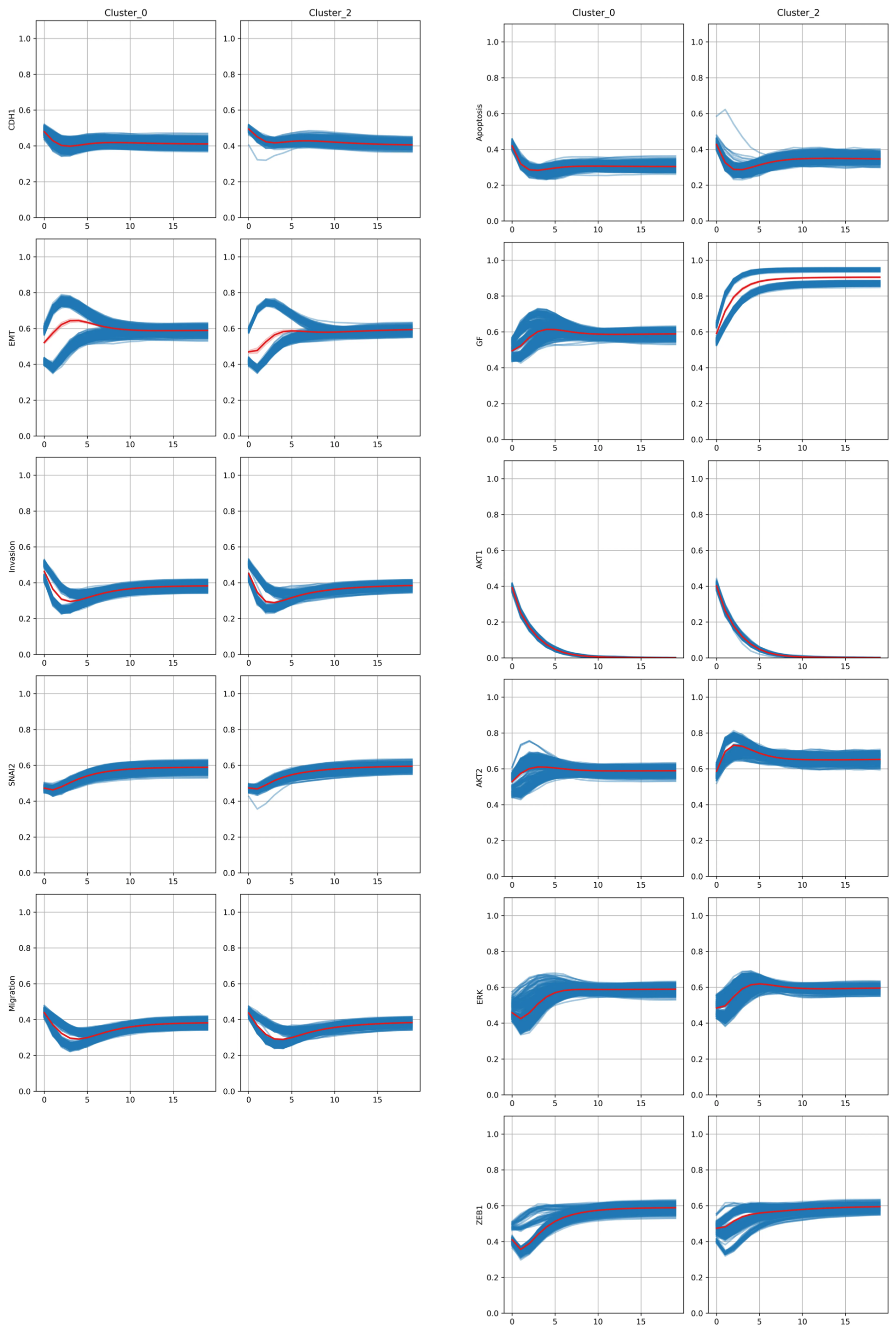


**Supplementary Figure S4.** Dynamics of selected node in two BN clusters from the invasion model network. Each plot shows the node activation probably of selected nodes (CDH1, EMT, Invasion, Apoptosis, GF, AKT1, AKT2, ERK, ZEB1) from the invasion model. The y-axis shows the level of node activation probability, and x-axis shows simulation timepoint from 0-20. The plot is splited into 2 according to the BNs clusters (cluster 0 and 2).

### 2 Supplementary Analysis

#### 2.1 Supplementary Analysis 1

In this part, we performed follow-up analysis on three cases with low Adjusted Rand Index (ARI) scores when comparing attractor-based and simulation-based clustering. The analysis was conducted on two models (Cortical and Human Gonadal), while the HCC1954 model was excluded due to its high number of reachable attractors (approximately 18,000 reachable attractors).

To investigate these differences, we utilized MaBoSS options to obtain states activation probability at the endpoint of simulation, exploring the likelihood of attractor states being reached. We visualized this comparison across four panels. The first panel displays the reachable attractor states in each model, with rows representing Boolean Networks (BNs) in the model ensemble and columns representing attractor states. Models are labeled with row colors corresponding to BN clusters that share attractor groups.

The second panel illustrates the states activation probability obtained from MaBoSS simulation, maintaining the same order of models and attractor states for direct comparison. The third panel presents the converted node activation probability calculated from the node activation probability. Finally, the fourth panel shows the Multidimensional Scaling (MDS) projection of model distances based on endpoint node activation probability, where each dot represents a model and colors indicate model clusters based on shared attractor groups.

We showed these plot with the network “Cortical Area Development_20862356” on the upper row, and the “Human Gonadal Sex Determination_26573569” on the bottom row.

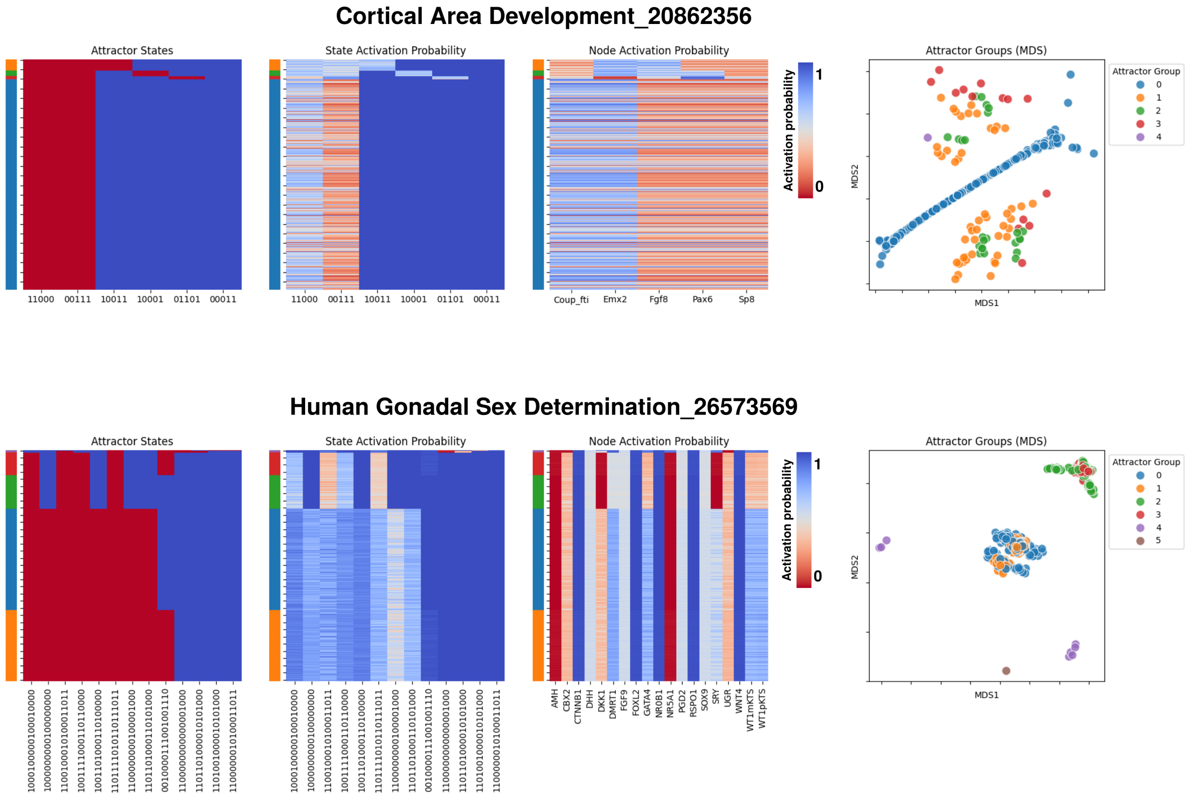


**Supplementary Figure S5.** The comparison of the calculated attractor, states activation probability and node activation probability for each BN in the model ensemble, along with each MDS projection of BN distance in each model ensemble. Each color of the dot in the project represent the attractor group of the BN
